# Complex structural variation is prevalent and highly pathogenic in pediatric solid tumors

**DOI:** 10.1101/2023.09.19.558241

**Authors:** Ianthe A.E.M. van Belzen, Marc van Tuil, Shashi Badloe, Alex Janse, Eugène T.P. Verwiel, Marcel Santoso, Sam de Vos, John Baker-Hernandez, Hindrik H.D. Kerstens, Nienke Solleveld-Westerink, Michael T. Meister, Jarno Drost, Marry M. van den Heuvel-Eibrink, Johannes H. M. Merks, Jan J. Molenaar, Weng Chuan Peng, Bastiaan B.J. Tops, Frank C.P. Holstege, Patrick Kemmeren, Jayne Y. Hehir-Kwa

**Affiliations:** Princess Máxima Center for Pediatric Oncology, Utrecht, The Netherlands; Department of Pathology, UMC Utrecht, Utrecht, The Netherlands; Oncode Institute, Utrecht, The Netherlands; UMCU-Wilhelmina Children’s Hospital - Child Health, Utrecht, The Netherlands; Division of Imaging and Oncology, UMC Utrecht and Utrecht University, Utrecht, The Netherlands; Department of Pharmaceutical Sciences, Utrecht University, Utrecht, The Netherlands; Center for Molecular Medicine, UMC Utrecht and Utrecht University, Utrecht, The Netherlands

**Keywords:** Complex structural variation, pediatric solid tumors, chromothripsis, chromoplexy, extrachromosomal DNA (ecDNA), whole genome sequencing (WGS)

## Abstract

**Background:** In pediatric cancer, structural variants (SVs) and copy number alterations can contribute to cancer initiation and progression, and hence aid diagnosis and treatment stratification. The few studies into complex rearrangements have found associations with tumor aggressiveness or poor outcome. Yet, their prevalence and biological relevance across pediatric solid tumors remains unknown.

**Results:** In a cohort of 120 primary tumors, we systematically characterized patterns of extrachromosomal DNA, chromoplexy and chromothripsis across five pediatric solid cancer types: neuroblastoma, Ewing sarcoma, Wilms tumor, hepatoblastoma and rhabdomyosarcoma. Complex SVs were identified in 56 tumors (47%) and different classes occurred across multiple cancer types. Recurrently mutated regions tend to be cancer-type specific and overlap with cancer genes, suggesting that selection contributes to shaping the SV landscape. In total, we identified potentially pathogenic complex SVs in 42 tumors that affect cancer driver genes or result in unfavorable chromosomal alterations. Half of which were known drivers, e.g. *MYCN* amplifications due to ecDNA and *EWSR1::FLI1* fusions due to chromoplexy. Recurrent novel candidate complex events include chromoplexy in *WT1* in Wilms tumors, focal chromothripsis with 1p loss in hepatoblastomas and complex *MDM2* amplifications in rhabdomyosarcomas.

**Conclusions:** Complex SVs are prevalent and pathogenic in pediatric solid tumors. They represent a type of genomic variation which currently remains unexplored. Moreover, carrying complex SVs seems to be associated with adverse clinical events. Our study highlights the potential for complex SVs to be incorporated in risk stratification or exploited for targeted treatments.

## Introduction

Structural variants (SVs) and copy number (CN) alterations occur frequently in pediatric cancer and can contribute to cancer initiation and progression. They can therefore also be leveraged for diagnosis and treatment stratification. Well-known examples include the *EWSR1::FLI1* fusion gene in Ewing sarcoma (EWS) [1], the *PAX3/7::FOXO1* fusion gene in fusion-positive rhabdomyosarcoma (FP-RMS) [2] and *MYCN* amplification in neuroblastoma (NBL)[3]. Recent studies have highlighted the potential clinical utility of considering the underlying mutational mechanisms of these driver alterations, as they may indicate differences in tumor aggressiveness. In EWS, patients with fusion genes arising from complex rearrangements have a worse prognosis than those with fusions resulting from simple reciprocal translocations [1]. Similarly, NBL patients with more complex structures of the *MYCN* focal amplification (amplicon) and co-amplification of oncogenes seem to form a higher risk group [4]. However, a systematic characterization of complex structural variation patterns across pediatric solid tumors is lacking.

Complex SVs are characterized by clusters of breakpoints reflecting the repair of multiple double strand DNA breaks that likely occurred at the same time. In recent years, many different complex SV classes have been distinguished based on patterns in sequencing data and hypotheses of how the DNA damage has occurred and was repaired. Examples include chromothripsis, chromoplexy, breakage-fusion bridge (BFB), local jumps, pyrgo, rigma and typhona [5, 6]. However, the criteria for categorization of complex SVs have been inconsistently used across studies, especially between cancer, developmental disease, and germline [7]. Recent mechanistic studies using single-cell sequencing have shown that many different complex SV patterns can arise from straightforward events during one cell division, such as chromatin-bridge breaking after telomere fusion resulting in focal chromothripsis-like patterns [8, 9]. Furthermore, the observed patterns of gains and losses could be explained by unequal division of genomic material and do not require complicated molecular mechanisms involving DNA synthesis [8, 9]. Considering these aspects, Zhang and Pellman (2022) conclude it might be difficult to distinguish the underlying mechanisms, such as BFB and chromothripsis, based on copy number patterns alone [8].

Alternatively, the umbrella term “chromoanagenesis” can be used to encompass a spectrum of complex SVs resulting in rearranged derivative chromosomes. Long-read and Hi-C data of germline genomes also support this diversity of rearrangements and categorization into chromothripsis-like and chromoplexy-like patterns [10]. Chromothripsis is characterized by DNA shattering and haphazard repair in which genomic fragments are randomly joined, resulting in many interleaved SVs and an oscillating CN pattern with losses [11]. On the other hand, chromoplexy is a CN balanced complex rearrangement characterized by translocations between multiple chromosomes. It can result in fusion genes such as *EWSR1::FLI1* and is thought to arise when DNA damage occurs during co-localization of chromosomes in transcription hubs [1]. Both chromothripsis and chromoplexy can result in derivative chromosomes, differing mainly in the presence or absence of large copy number changes [10, 12].

In contrast, extrachromosomal DNA (ecDNA) fragments can result in high-level amplifications of oncogenes. A well-studied example includes *MYCN* amplification in NBL: an initiating event excises the oncogene after which it is amplified as circular ecDNA fragment [4, 13]. The ecDNA can undergo further rearrangements and either remain as an extrachromosomal fragment or integrate back into the genome as a homogeneous staining region [14]. The initiating event usually has characteristics of chromothripsis or BFB and can also include additional genomic loci such as cancer genes or distal enhancers [4, 13, 14]. This can give rise to a great diversity of ecDNA amplicon structures, and research is ongoing into associations with treatment resistance, tumor aggressiveness and patient outcome [14–16].

Here, we systematically characterized complex SV patterns across pediatric solid tumors and identified candidate pathogenic events that likely contribute to tumorigenesis. To study complex SVs agnostic to classes or types, we first detected SV clusters representing complex events and subsequently categorized them into chromothripsis, chromoplexy or ecDNA/amplicons. We found that complex SVs occur in approximately half of our cohort (56 of 120 tumors) and the same complex SV classes occur across multiple cancer types, indicating similarities in mutational mechanisms. Furthermore, recurrently altered genomic regions often overlap with cancer genes and are mostly cancer-type specific, which suggests that selection is involved in shaping the observed complex SV landscape. In 75% of tumors carrying a complex SV (42 tumors), we identified candidate pathogenic complex events that affect cancer driver genes or result in unfavorable chromosomal alterations associated with poor prognosis. In addition, patients whose tumor carried a complex SV involving a cancer driver gene experienced a clinical event twice as often compared to patients without. In conclusion, our results indicate that complex SVs are highly pathogenic in pediatric solid tumors and that the clinical implications of complex SVs warrants further study.

## Results

### Patterns of complex structural variation across pediatric solid tumors

To investigate pan cancer patterns of chromothripsis, chromoplexy and ecDNA, we analyzed somatic structural variants in a cohort of 120 patients across five types of pediatric solid tumors. Paired tumor-normal WGS data was generated from primary tumors of patients diagnosed with neuroblastoma (NBL, n=39), Ewing sarcoma (EWS, n=22), Wilms tumor (WT, n=34), hepatoblastoma (HBL, n=7) and rhabdomyosarcoma (RMS, n=18). Overall, these tumor genomes have an anticipated low tumor mutation burden (TMB) of single nucleotide variants (SNVs) and indels (median 0.32 per Mb) and fraction of genome altered by copy number alterations (FGA, median 12.6%) (Figure 1a) compared to adult cancers [17]. Additionally, we identified a median of 14 somatic SVs with >0.1 tumor allele fraction (Figure 1a). To infer complex events, we identified clusters of SV breakpoints within a 5 megabase pair (Mbp) interval and categorized them into different types of complex events based on a combination of SV and CN features (Figure 1b).

**Figure 1:**
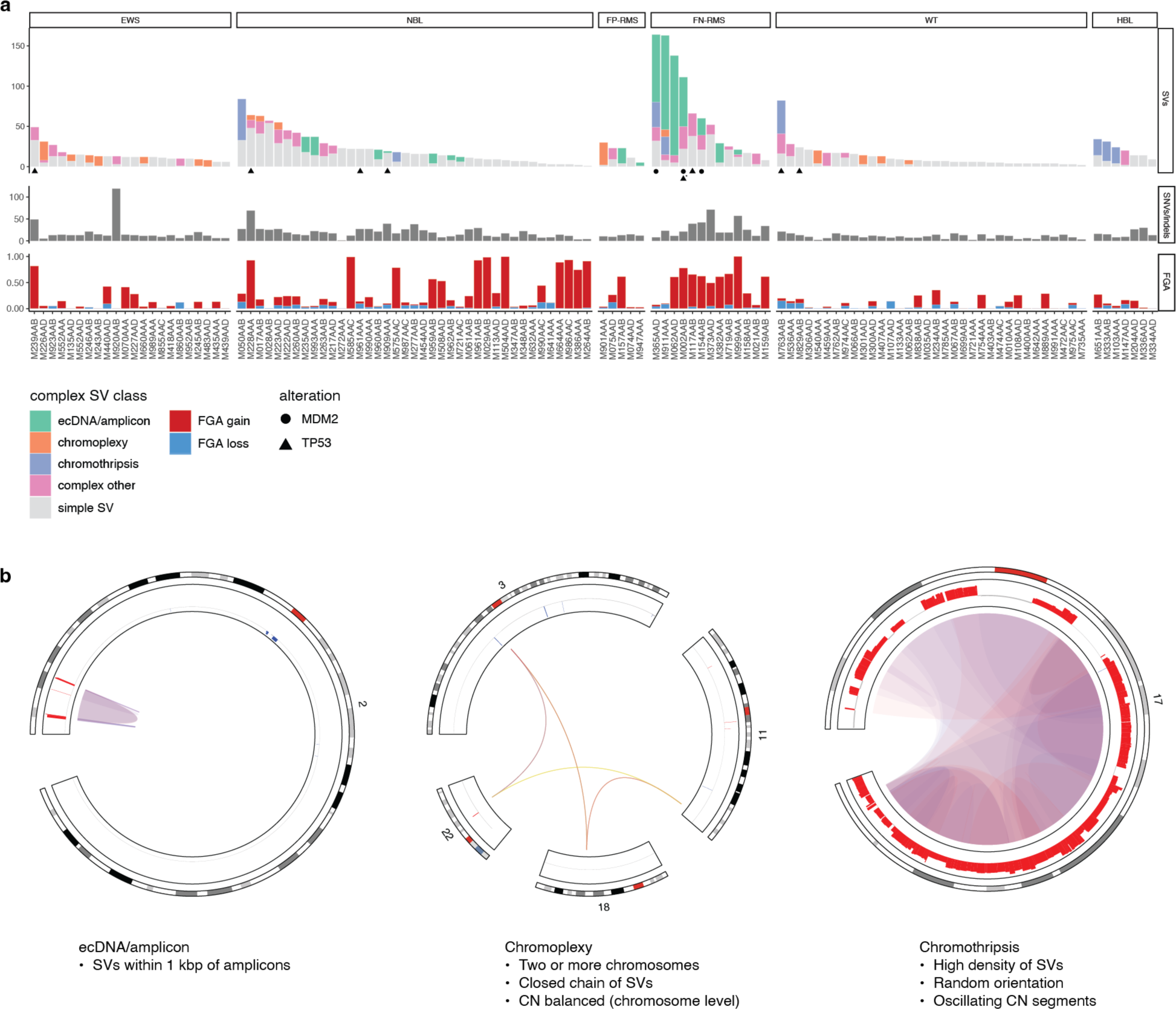
Occurrence of complex SVs in pediatric solid tumors. **a** Number of SVs per tumor colored by complex SV class (top), number of nonsynonymous SNV/indels (middle), and fraction of genome altered by copy number gain or loss (bottom) across pediatric solid tumors from left to right: Ewing sarcoma (EWS), neuroblastoma (NBL), fusion-positive and fusion-negative rhabdomyosarcoma (FP-RMS, FN-RMS), Wilms tumor (WT), and hepatoblastoma (HBL). Symbols denote genomic instability mutations: TP53 disruption (triangle), MDM2 amplification (circle). * M002AAB has a germline TP53 alteration. **b** Characteristics of complex SV classes and circos plots of examples, left to right: extrachromosomal DNA (ecDNA)/amplicon-type (patient M721AAC), chromoplexy (patient M135AAD) and chromothripsis (patient M050AAB). SV clusters are categorized based on the following core characteristics. ecDNA/amplicon: SVs with breakpoints within 1 kilobase pair (kbp) of amplicons. Chromoplexy: SVs form a closed cycle and connect multiple chromosomes via copy number (CN) balanced interchromosomal breakpoints. Chromothripsis: footprints with oscillating CN segments and at least 10 overlapping SVs of mixed types, indicating randomly joined fragments. SV clusters not complying to these criteria, were categorized as “complex other”. See Methods for details.

Applying this SV clustering and categorization approach, we identified complex SVs in 56 tumors (47%), across all five cancer types (Figure 1, Table 1). Most tumors carry a single complex event, but those with mutations in *TP53* or *MDM2* carry multiple complex SV events (median 3.5) and have a higher TMB (median 0.80 vs 0.32, p<0.01) and FGA (41% vs 12%, p<0.01). However, not all tumors with complex SVs have a mutation in *TP53 or MDM2* illustrating that it is not a prerequisite for complex rearrangements to occur or to be tolerated. Each complex SV class was identified in at least two of the five solid cancer types. In total, chromoplexy was identified in 16 tumors (EWS, WT, RMS, and NBL), ecDNA/amplicon in 16 tumors (NBL and RMS), and chromothripsis in 8 tumors (NBL, WT, HBL, and RMS), indicating that complex SVs occur in many pediatric solid tumors.

Chromoplexy was the most prevalent in Ewing sarcoma (EWS), it was detected in seven tumors as the underlying mechanism generating the pathognomonic driver fusions with *EWSR1.* In four other EWS, we also identified complex SVs underlying the driver fusions. However, these were categorized as “other” as they either did not form a closed cycle or included an unbalanced translocation and therefore did not pass all criteria for canonical chromoplexy. Chromoplexy was also detected in RMS (n=2), NBL (n=3) and WT (n=4), making it the most widespread class of complex SV. Many of these events will remain undetected by exome sequencing given their CN balanced nature, stressing the importance of performing WGS as part of the molecular diagnostic process.

Amplicon-overlapping complex SV clusters were identified in NBL and RMS. In seven of nine NBLs with *MYCN* amplification, we detected overlapping complex SV clusters that likely reflect ecDNAs and typically consist of interleaved SVs of mixed types, sizes and variant allele fractions, as expected [4, 13]. In addition, we identified SVs connecting multiple amplified loci that indicate these are part of the same ecDNA construct (Table 2). Apart from *MYCN* amplification in NBLs, we also identified complex SVs overlapping amplicons in nine RMS tumors. Although their genomic locations differ, the underlying rearrangements of these complex events resemble those of the NBLs, including amplified loci on different chromosomes that have become physically linked. This suggests that ecDNA is a more widespread phenomenon than *MYCN* amplifications in neuroblastoma alone.

Chromothripsis-like complex SVs were identified across multiple cancer types in eight tumors. Full-chromosome chromothripsis was rare in our cohort. We detected it in only two NBLs: M050AAB has an additional chromothriptic copy of chr17 and M575AAC has subclonal chromothripsis of chr15. However, focal chromothripsis was identified more often, namely in HBL (n=3), WT (n=1), and RMS (n=2), and resulted in either loss of the remainder of the chromosome arm (n=4) or novel derivative chromosomes (n=2). The SV and CN patterns of these complex events are consistent with the proposed mechanism of focal chromothripsis arising from chromatin bridge breaking during a single cell division [9]. Furthermore, the presence of chromothripsis did not require *TP53* or *MDM2* to be altered. One WT with focal chromothripsis had biallelic *TP53* disruption, but we did not identify alterations in *TP53* or *MDM2* in the other seven tumors with chromothripsis. This substantiates that chromothripsis can result from a single event and does not necessarily indicate ongoing genomic instability due to inactivation of the P53 pathway.

Overall, the characteristic patterns of ecDNA/amplicons, chromoplexy and chromothripsis were commonly observed across multiple solid cancer types. In line with previous work, we identified chromoplexy underlying fusion genes in EWS and ecDNA underlying *MYCN* amplifications in NBL. However, we also identified similar complex SVs in other cancer types. Therefore, we further investigated pan cancer patterns of complex SVs and assessed their potential contribution to tumorigenesis.

### Complex SV hotspots point to potentially pathogenic events and not genome fragility

To analyze whether complex SVs affect the same genomic regions across different tumors, we conducted a systematic, genome-wide analysis and identified genomic regions that have SV breakpoints in more than 2% of our cohort (n≥2 Figure 2, Table 3). In total 13 hotspot regions were affected by complex SVs in three or more tumors. Among these hotspots were regions mostly comprised of SVs underlying known driver alterations, namely with *FOXO1, EWSR1, FLI1, MYCN,* and *ALK*. In addition, novel complex SV hotspots outside of the known driver events were identified on chromosomes 1, 11, 12, 16 and 17. These include candidate regions of interest that harbor cancer genes, such as *WT1* and *MDM2,* or where chromosomal alterations have previously been associated with poor outcome, such as chr1p [3, 18, 19]. Furthermore, the breakpoints comprising the complex hotspots often originate from the same cancer type (Figure 2).

**Figure 2:**
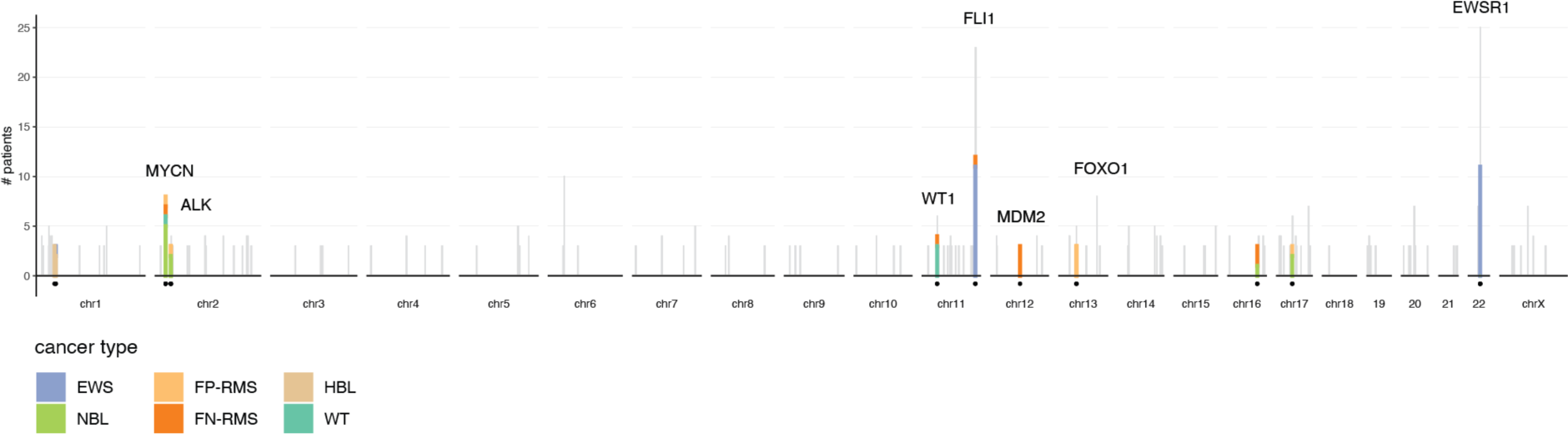
Complex SV hotspots contain cancer driver genes. Genome-wide overview of recurrently altered genomic regions with SV breakpoints from three or more tumors (gray bars). Complex SV hotspots containing complex SV breakpoints from three or more tumors are highlighted (circles, thick lines), colored according to their cancer type and annotated with overlapping cancer driver genes.

To assess whether the hotspots reflect genome sequence propensity for rearrangement, we checked if they were enriched in repetitive elements. Whilst a fraction of all SVs is likely repeat-mediated, with both start and end breakpoints nearby repeat elements of the same class, these are less often part of a complex SV cluster and instead occur as simple SVs (7.6% vs 27%, p<0.01). When only considering complex SVs in hotspots, an even lower proportion is possibly repeat-mediated (4%). Consistent with this, the complex hotspots are depleted of repeat-mediated SVs (6% vs 18%, p<0.01) compared to the remainder of the genome, regardless if they are simple or complex, suggesting that the observed recurrence is not likely due to genome fragility.

Alternatively, we hypothesized that selection plays a role in the formation of complex SV hotspots. Large-scale SVs are highly dysregulating and therefore unlikely to be selectively neutral, so the observed recurrence can indicate that these alterations provide a competitive advantage to the tumor cells [20]. Complex SV hotspots are significantly enriched for SVs with breakpoints inside (46% vs 5.9%, p<0.01) or nearby (93% vs 34%, p<0.01) known pediatric cancer genes. This does not solely arise from SVs being either complex or simple. Comparing complex to simple SVs shows that they have only slightly more breakpoints nearby pediatric cancer genes (43% and 36%, p<0.05) and similar fractions inside the gene body (9.4% and 9.3%, not significant). However, there are marked differences between complex SV classes: 26% of chromoplexy breakpoints are located inside gene bodies compared to just 5.7% and 4.5% for ecDNA/amplicons and chromothripsis, respectively. Since chromoplexy is thought to involve transcription hubs, the genes with chromoplexy breakpoints are likely highly expressed and important for cell function. For ecDNA/amplicons, the breakpoints tend to flank the genes instead of residing in gene bodies, but the amplified region is usually confined to known oncogenes, also indicating specificity. In contrast, chromothripsis events tend to cause more widespread disruptions and have more breakpoints overall, of which some can intersect genes by chance. Since chromothripsis is expected to arise from random breakage, recurrent events are of particular interest. Taken together, most hotspots either reflect cancer driver events known to occur in these pediatric cancers, or harbor other relevant cancer genes or chromosomal alterations, suggesting that these recurrent complex SVs are potentially pathogenic events.

### Complex SVs provide insights into underlying genomic rearrangement of driver alterations

To identify potentially pathogenic complex SVs, we selected events that affect cancer-type specific driver genes or result in unfavorable chromosomal alterations that have been associated with poor outcome or high risk (Table 1, Figure 3). In total, we identified candidate complex events in 42 tumors (75% of tumors with complex SVs). Half of which are drivers known to arise from complex rearrangements (n=21), namely *EWSR1::FLI* fusion genes, *MYCN* amplifications and *PAX3/7::FOXO1* fusion genes. When these alterations are identified, they are considered to be the main driver events for these cancer types. Consistent with this, these events were the only complex SV found in most of these tumors (n=19 of 21, Figure 3). Furthermore, even though these complex SVs give rise to recurrent driver events, their underlying genomic rearrangements show a large variation among the tumors.

**Figure 3:**
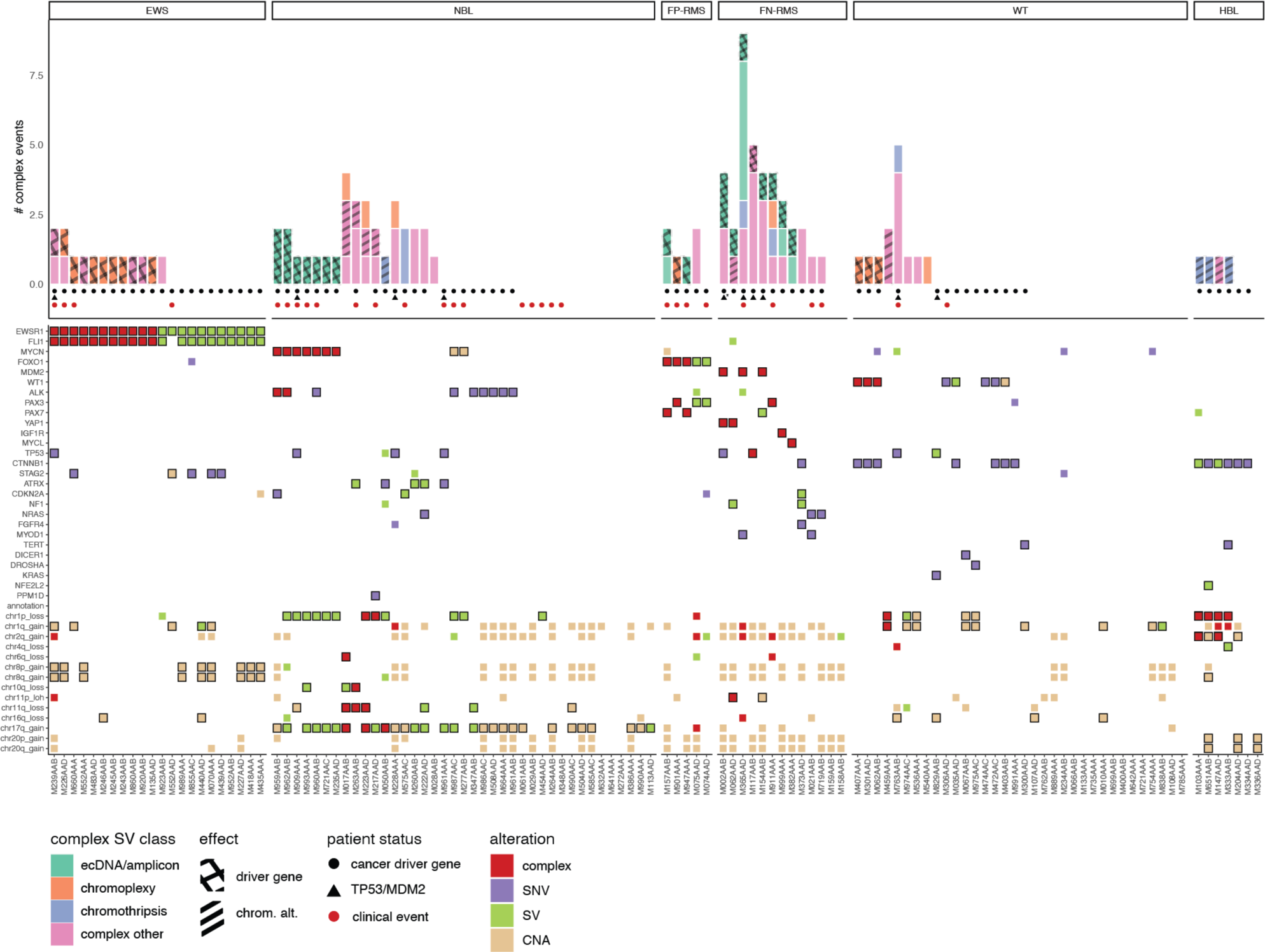
Complex SVs affect known cancer driver genes and chromosomal alterations. Number of complex events (top) per tumor colored by complex SV class and filled with patterns indicating their effect: cancer driver gene (cross) or unfavorable chromosomal alteration (lines). Patients are annotated by whether the tumors carry a clinically relevant driver alteration (circle) or an alteration in TP53 or MDM2 (triangle), as well as whether they experienced a clinical event (progression, relapse or death (red circle)). * M002AAB has a germline TP53 alteration. Mutation status per tumor (bottom) across selected genes and genomic regions that are relevant in at least one of the included cancer types. Alterations are colored by type: complex (red), SV (green), SNV/indel (purple) or copy number alteration (CNA, beige). Alterations of known relevance in that cancer type are highlighted by a black border.

The SVs underlying the *EWSR1::FLI* fusion genes all have breakpoints in the *EWSR1* and *FLI1* hotspots, whether they are simple (n=11) or complex (n=11). Although some SVs from non-EWS tumors also occur in this region, they do not have breakpoints that map inside the genes and therefore probably do not have the same functional effect. Whilst the simple reciprocal translocations between *EWSR1* and *FLI1* only affect these specific genes and result in the balanced t(11;22), the complex rearrangements can additionally affect other cancer genes and result in different derivative chromosomes (Table 1).

The structure of the *MYCN* amplicons differs across NBL tumors, sometimes involving additional enhancers or cancer genes in the resulting ecDNA (Table 2). For example, some tumors display amplification of a broad consecutive region likely including the downstream e4 enhancer as it extends beyond 16.4 Mbp [4]. In contrast, other ecDNAs are comprised of multiple separate regions, such as upstream loci or full cancer genes. For example, we identified co-amplification of *MYCN* and *ALK*, and of *FBXO8* and *CEP44* which have both been associated with poor outcome [4, 21].

For the five fusion-positive RMS tumors, we found profound differences in the underlying genomic alterations of their driver fusion genes (Figure S1). Despite recurrent breakpoints in specific locations of the *PAX3/7* and *FOXO1* genes, the underlying structural rearrangements are different. Two tumors carry *PAX7::FOXO1* fusion genes and in both cases, we identified ecDNA/amplicon-type complex SVs resulting in amplification of the driver fusion. In addition, tumor M157AAB carries a *MYCN* amplification and SV patterns indicate it is likely part of the same ecDNA as its *PAX7::FOXO1* fusion gene. In contrast, for two of three tumors with *PAX3::FOXO1*, the underlying rearrangements are unbalanced reciprocal translocations and for one it originated through chromoplexy. Furthermore, fluorescence *in situ* hybridization using *FOXO1* break-apart probes supported that these five fusion-positive RMS have different underlying rearrangements (Figure S2). Overall, the individual differences we observed in driver alterations between tumors provide opportunities for applying precision medicine approaches when treating high-risk cancers.

### Most complex SVs confer an advantage to the tumor

In addition to drivers known to arise from complex rearrangements, we identified candidate complex events in 21 tumors to investigate further (Table 1). This includes complex SVs affecting driver genes usually mutated by SNVs/indels, as well as complex rearrangements resulting in gains or losses relevant to the cancer type. A subset of these candidate complex events has breakpoints in the complex SV hotspots on chromosomes 1, 11, and 12, they reflect cancer-type specific recurrent events. No other driver alterations have been identified in six tumors carrying candidate complex events (Figure 3, Table 4), increasing the likelihood that these complex SVs are pathogenic. Also, patterns in the variant allele fractions and CNs indicate many complex events are clonally present (Table 1, Table 5). In tumors with known complex drivers, we observed few additional complex SVs. Similarly, in 10 of 21 tumors with a candidate complex event, it was the only complex SV we identified.

Loss of the tumor suppressor gene *WT1* is an important cancer driver event in Wilms tumor (WT) [22]. In three WTs, we identified chromoplexy breakpoints that reside inside *WT1* and result in rearranged chromosomes and substantial disruption of the gene (Figure S3). The complex SV was fully CN balanced for tumor M062AAB, making it impossible to detect with exome sequencing or targeted assays. For the remaining two tumors, it is accompanied by focal deletions at the breakpoints, but the resulting derivative chromosomes would go undetected without the use of WGS. Furthermore, these chromoplexy breakpoints are located in the complex SV hotspot on chr11 which also harbors breakpoints from simple SVs affecting *WT1* in two other WTs (Table 3). In all of these five WTs, the SVs are clonally present, and the tumors do not carry SNVs or indels in *WT1.* This further stresses the importance of analyzing SVs in WTs to detect all pathogenic events.

Focal chromothripsis resulting in loss of chr1p was detected in three of the seven hepatoblastomas (HBLs). They all have breakpoints located in a hotspot on chr1 and loss of the adjacent region of chr1p (Figure 4, Table 5), indicating that this focal chromothripsis represents a recurrent event in HBL. We identified a shared lost region on chr1p (chr1:1-34,847,815) and in two tumors the complex SV also resulted in either chr1q or chr2q gain. All three of these chromosomal alterations have been associated with tumor aggressiveness and/or poor prognosis [18, 23]. The remaining fourth complex SV we identified in a HBL tumor also results in 1p loss, 1q gain, and 2q gain (Figure S4). Moreover, these instances of complex SVs are the only complex events in these tumors and seem to be clonally present. In conclusion, we identified four complex events in HBLs that all result in unfavorable chromosomal alterations, indicating that they likely provide proliferative advantages.

**Figure 4:**
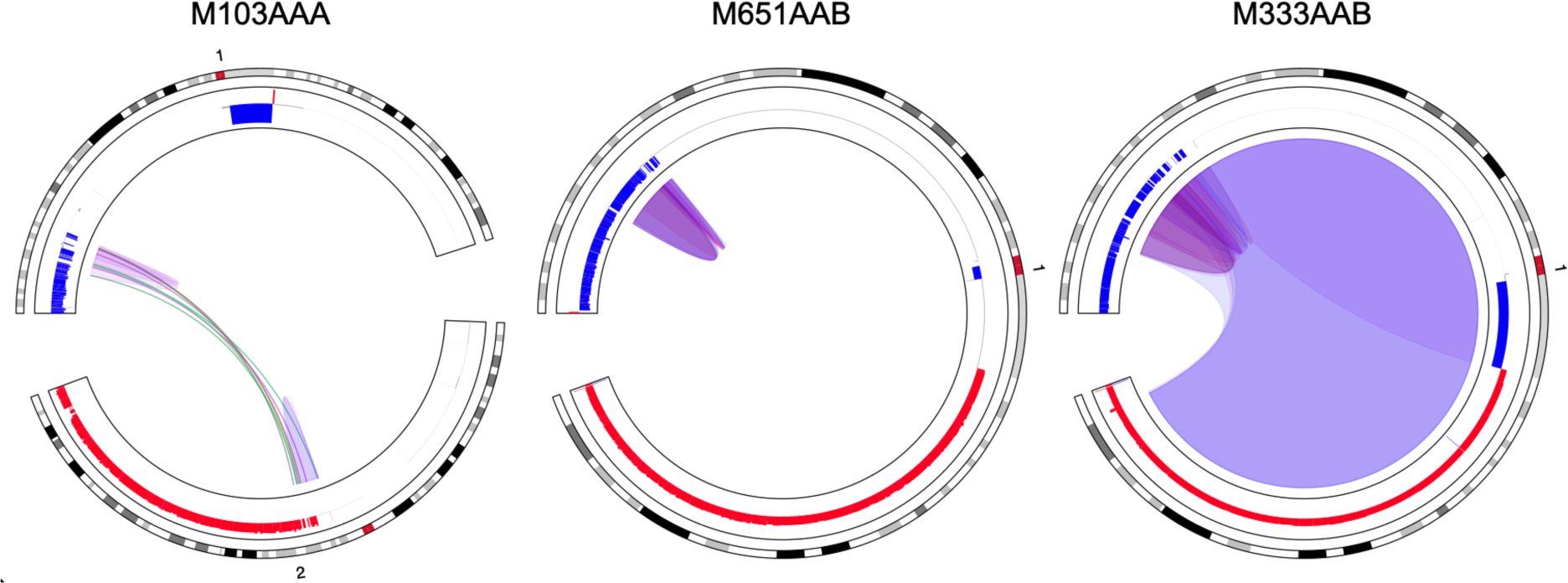
Focal chromothripsis in hepatoblastoma. Recurrent focal chromothripsis was identified in three hepatoblastomas, from left to right: tumors from patients M103AAA, M651AAB, and M333AAB. All three examples have breakpoints in the complex SV hotspot on chromosome 1 (chr1:33517560-35493723) (Table 3) and a recurrently lost region on chr1p: chr1:1-34847815 (Table 5).

In five neuroblastomas (NBLs) we identified complex SVs resulting in chromosomal alterations that have been associated with poor outcome, such as chr1p loss, chr11q loss, and chr17q gain [3, 24, 25] (Figure S5). These candidate complex SVs are not recurrent at the breakpoint level but result in recurrent CN changes (Figure 3). In tumor M050AAB we identified chromothripsis of chr17 present as an additional derivative chromosome, effectively leading to chr17q gain. The other four complex candidate events consist of multiple unbalanced translocations that result in one or more unfavorable chromosomal alterations at the same time (Figure S5). For example, tumor M263AAB carries a complex event connecting multiple unbalanced translocations resulting in chr10q loss, chr11q loss, and chr17q gain. This shows that single instances of complex SVs can have a large impact on the tumor genome, and that markers of poor prognosis can be interdependent and physically linked.

Amplification of oncogenes via ecDNA can be a potent cancer driver event as illustrated by *MYCN* amplification and the *PAX7::FOXO1* fusion genes. Furthermore, we identified potentially pathogenic ecDNA/amplicon-type complex SVs in seven fusion-negative RMS tumors, resulting in amplification of *MDM2*, *MYCL*, *IGF1R1,* and *YAP1* (Figure S6). On chr12 we identified a hotspot overlapping with the *MDM2* oncogene that consists of ecDNA-like breakpoints from three RMS tumors and likely reflects a pathogenic *MDM2* amplification (Figure S6). This includes breakpoints from *MYOD1*-mutated tumor M365AAD, where we observed a pattern of interconnected focal gains that did not meet the amplicon threshold but nevertheless suggests the presence of multiple subclonal ecDNAs (Figure S7). Of special interest is the ecDNA identified in tumor M911AAA resulting in the rare fusion gene *PAX3::WWRT1* (Figure S8). Previously, we found that the transcription profile of this particular tumor clustered with canonical fusion-positive RMS tumors which form a high-risk subgroup [26]. Moreover, for three of these seven fusion-negative RMS with ecDNA/amplicons resulting in oncogene amplification, we found no alterations in cancer driver genes outside of these complex events (Figure 3).

In conclusion, for 42 of 56 tumors carrying a complex SV, we identified potentially pathogenic candidate complex events affecting cancer-type specific driver genes or unfavorable chromosomal alterations. Furthermore, twice as many patients whose tumors have a complex SV affecting a driver gene experienced an adverse clinical event such as progression, relapse, or death, compared to those without such a complex event (41% vs 19%, p<0.05, odds ratio 2.8, Figure 3). This cohort-wide observation is consistent with earlier findings of the relationship between carrying complex SVs and poor outcome which were based on examples in specific cancer types or complex SV classes [4, 15, 27, 28]. Taken together, our results show that complex SVs tend to be pathogenic events that play an important role in tumorigenesis and are likely indicators of tumor aggressiveness.

## Discussion

Structural variants are important drivers of pediatric cancer, but the prevalence and biological relevance of complex rearrangements has remained elusive. We systematically characterized patterns of extrachromosomal DNA, chromoplexy and chromothripsis in a cohort of 120 primary tumors across five types of pediatric solid tumors. Complex SVs are prevalent, with almost half of all tumors having at least one complex event and instances of complex SV classes occurring in multiple cancer types. Hotspot regions that are recurrently altered by complex SVs often overlap with cancer genes and tend to contain breakpoints from the same cancer type, suggesting that selection pressures play a role in shaping the SV landscape. Overall, we identified candidate pathogenic complex SVs in 35% of all tumors that affect genes or chromosomal alterations known to be relevant in these cancer types. Half of which were known drivers with underlying complex rearrangements, e.g. *EWSR1::FL1* fusion genes and *MYCN* amplifications. In addition, we identified complex SVs affecting genes or regions previously associated with poor outcome, such as *MDM2* amplification in fusion-negative RMS due to ecDNA/amplicons, and chr1p loss in HBLs due to focal chromothripsis. In conclusion, complex SVs are prevalent and highly pathogenic in pediatric solid tumors.

Despite the relatively low mutation burden of pediatric cancer, complex SVs occurred frequently including in tumors with few copy number (CN) alterations and SVs, suggesting an overall absence of (ongoing) genomic instability. Furthermore, many tumors carry only a single complex event with variant allele fractions and CN patterns indicating clonal presence. In contrast, adult cancers have higher mutation burdens and are usually heavily altered by both simple and complex rearrangements, with chromothripsis occurring in ∼30% of tumors and chromoplexy in ∼18% [5, 29]. Disruption of *TP53* has been linked to a higher prevalence of chromothripsis in both pediatric and adult cancers but is not a prerequisite for complex SVs to occur [17, 29]. This further supports the notion that complex SVs do not reflect generic genomic instability but rather reflect one-off events with specific mutational mechanisms. However, to compare complex SV patterns between cancer types requires striking a balance between consistent definitions and tailored detection methods. Since complex SVs were first discovered in adult cancers, criteria for their detection and classification were made with high background mutation rates in mind [11]. To accommodate observations of complex events in germline genomes, some studies utilized relaxed criteria, which makes it difficult to compare outcomes and can result in obfuscation of complex SV classes [7]. Although dedicated detection tools such as Shatterseek [29] can provide uniformity, we found that this statistical approach was less suited to our dataset of pediatric cancer genomes with very few breakpoints. Therefore, we identified complex events agnostic to SV class by detecting clusters of SV breakpoints and performed categorization into the different complex classes afterwards. This allowed for a systematic investigation of complex events across pediatric cancer genomes.

Among pediatric solid tumors, we observed distinct complex SV patterns likely arising from an interplay between similar mutational mechanisms and different selection pressures. Although differences in 3D genome structure can also contribute to the observed cancer-type specific patterns [20], the fact that often only a single complex SV occurs in a pediatric solid tumor genome and it regularly affects known driver genes suggests that both negative and positive selection contribute to the observed complex SV landscape. For EWS and NBL, it has been established that complex rearrangements play a role as cancer drivers [1]. In EWS, we identified complex rearrangements underlying half of the driver fusions but detected no chromothripsis or ecDNA, which is in line with previous studies [27, 30]. In NBL, we can distinguish two subgroups with different complex SVs and a third group largely devoid of complex SVs: 1) *MYCN* amplified (MNA) tumors, 2) tumors carrying chromothripsis, chromoplexy and related events, and 3) hyperdiploid NBLs which are generally classified as lower risk [24]. For all tumors with *MYCN* amplification due to ecDNA, it was the only complex event present in the tumor genome, stressing its importance as a tumor driver. Moreover, we found differences in amplicon structure such as inclusion of enhancers or co-amplification of additional cancer genes, some of which previously have been associated with poor outcome [4, 21]. The second subgroup of tumors represent high-risk non-MNA tumors that carry complex SVs resulting in derivative chromosomes. Our observations fit the three mutational scenarios recently proposed by Rodriguez-Fos *et. al.*, namely 1) reactive oxygen species and replication stress, 2) homologous recombination repair deficiency, and 3) chromosome missegregation [31]. However, to our knowledge, we are the first to report complex SVs resulting in multiple unfavorable chromosomal alterations such as chr1p loss, chr11p loss, and chr17q gain [3, 24]. Considering the physical linkage of chromosomal alterations could provide additional insights and ultimately improve clinical decision making.

Extrachromosomal DNA (ecDNA) has been associated with oncogene amplifications and poor outcome in multiple cancer types [4, 13, 15]. The impact of ecDNA on tumor biology is profound, with its structure leading to severe dysregulation of genes and non-linear inheritance contributing to intratumor heterogeneity [14]. The persistence of ecDNAs without centromeres is likely due to positive selection, as indicated by computational models [14, 32], cancer-type specific differences in what genes are amplified [14, 15], as well as changes in the abundance of ecDNAs as response to drug treatment [16]. Although oncogene amplifications have been reported in RMS [2, 33], the prevalence of ecDNA/amplicon-type complex events in RMS was unanticipated. For three patients where we identified *PAX* fusions arising from ecDNA in the primary tumor, we verified the presence of the same ecDNA breakpoints at relapse (M911AAA and M947AAA) or in organoid culture (M157AAB) (Supplementary figure S9) [34]. This indicates that the ecDNAs are maintained and likely provide a selective advantage. Furthermore, M947AAA’s tumor acquired *MYCN* amplification during relapse, where it is linked to its *PAX7::FOXO1* fusion in a single ecDNA/amplicon-type complex SV. This resulted in a similar construct as present in tumor samples from M157AAB. Complex ecDNA structures like this can arise from clustering together of separate ecDNAs in hubs, followed by recombination into a single structure [14]. Although *MYCN* amplification is common in fusion-positive RMS [35], to our knowledge, co-amplification of *MYCN* and *PAX7::FOXO1* on a single amplicon structure has not been reported previously. Since we identified ecDNAs containing *PAX7, FOXO1* and *MYCN* in tumors from two different patients, it does not seem to be an incidental finding but rather reflect difficulties in detecting these events without WGS. Furthermore, there have been conflicting reports regarding the association of *MYCN* amplification with outcome in fusion-positive RMS [35], but the underlying rearrangements were not considered in these studies. In addition to driver fusions, we identified complex SVs underlying oncogene amplification in fusion-negative RMS, e.g. *MDM2*, *YAP1* and *IGF1R*, which could provide leads for targeted therapies [23]. Moreover, previous studies found that *MDM2* was among the oncogenes most often amplified as circular ecDNA across cancer types [15, 36]. Not only did circular amplicons achieve higher CNs, also the oncogene expression was higher corrected for CN due to increased chromatin accessibility and possible enhancer rewiring [15]. Since short-read sequencing is limited in its ability to fully resolve amplicon structures, this highlights a need for additional technologies such as long-read sequencing or CIRCLE-seq [37]. Even then, ecDNAs can be heterogeneous within a tumor cell population, and are able to transition between extrachromosomal states and integration into chromosomes [14, 16]. This flexible nature also contributes to their pathogenicity. Detection of these highly amplified complex SVs is a first step towards assessing their clinical relevance.

By considering complex SVs, we identified potentially pathogenic variants in WT and HBL. For example, in three Wilms tumors lacking a driver SNV, chromoplexy breakpoints disrupted *WT1*. WTs are usually regarded as “genomically quiet” since they carry few recurrent genetic alterations [22]. However, we found that WT genomes can contain “hidden” impactful variation, such as novel derivative chromosomes, that can be uncovered and studied using WGS. Similarly, HBLs carry few mutations and have a paucity of known genetic drivers outside of *CTNNB1*-activating mutations [38]. Yet four of the seven tumors recurrently altered chromosome 1p via a complex SV. Taken together, we identified known examples of complex cancer driver events, as well as additional likely pathogenic complex SVs in cancer types where it was not anticipated. Since we also observed that patients experienced a clinical event twice as often when their tumor carried a complex SV affecting a cancer driver gene, the clinical implications of complex SVs warrant further study.

## Conclusions

Complex SVs commonly occur in pediatric solid tumors and our findings highlight the importance of analyzing them as candidate pathogenic events. Not only do complex SVs give rise to known driver alterations such as fusion genes and oncogene amplifications, but they can also affect cancer genes usually altered by SNVs/indels or result in relevant CN gains and losses. Interpreting complex SVs as events that can have multiple effects on genome structure at the same time enables more refined functional analysis of genetic alterations. Moreover, CN balanced complex SVs are largely an unexplored type of SV and yield promising new candidates in tumors without known driver alterations. Finally, carrying complex SVs has previously been associated with tumor aggressiveness and poor outcome in specific cancer types, and our findings support this agnostic to the type of tumor. Complex SVs affect a significant proportion of pediatric solid tumors, and our findings warrant further research into their role in cancer etiology and progression.

## Materials and Methods

### Cohort selection and sequencing

To characterize complex structural variation patterns across pediatric solid tumors, we selected patients diagnosed with Ewing sarcoma, neuroblastoma, rhabdomyosarcoma, Wilms tumor and hepatoblastoma for which primary tumor material was included in the Máxima biobank, subject to informed consent [39]. Patients were eligible when whole-genome sequencing (WGS) data of sufficient quality was available from matching tumor-normal samples taken within 150 days of diagnosis and the variant calling steps were successfully completed. Sequencing library preparation and data pre-processing, including alignment and quality control, was done via the institute’s standardized pipelines and guidelines as described before [39–41]. In summary, high quality WGS samples were selected requiring a minimum median coverage of 27x for normal samples and 81x for tumor samples. We also included two samples with lower coverage of the tumor (PMABM000DEP with 60x, PMABM000DIX with 80x) that have been successfully analyzed previously [40]. In total, for 120 patients WGS data was available with a median coverage of 37x for the normal and 107x for the tumor samples (Table 6).

### Single nucleotide variants and indels

Somatic single nucleotide variants (SNVs) and indels were identified using Mutect2 from GATK 4.1 [42] and annotated by variant effect predictor (VEP) (version 104) [43]. First, we filtered high-confidence variants that have tumor allele fraction > 0.1 and are located on chromosomes 1-22 and X, excluding ENCODE Blacklist poor mappability/high complexity regions [44]. Second, to select likely pathogenic variants, we filtered on VEP impact moderate or high and removed variants predicted as benign/tolerated by PolyPhen/SIFT unless they were present in COSMIC [45].

The tumor mutation burden (TMB) was defined as the number of nonsynonymous somatic SNVs and indels per megabase. This encompasses the SNVs/indels in protein coding genes on chromosomes 1-22 that also passed the previous filtering steps. For the denominator, we used ∼41 megabase pairs (Mbp) corresponding to the number of coding sequence bases in protein coding genes.

### Copy number alterations

Copy number (CN) alteration data was generated by the GATK4 pipeline following their recommended best practices [42]. Across all analyses, we distinguished four CN call states: gain, loss, loss of heterozygosity (LOH) and neutral (no change). For gain and loss we required at least +/- 0.2 copy ratio log2 fold change (cr l2fc), and for LOH less than 0.4 minor allele fraction (MAF) with absence of gain/loss. The remainder was regarded as neutral. As a proxy for assessing the CN stability across a genomic region, we used the percentage of sequence within 33% of the mean CN for gain/loss and between -0.1 and 0.1 cr l2fc for neutral. Genomic regions with at least 70% of sequence near the target value were regarded as stable.

The fraction of the genome altered by copy number alterations (FGA) was calculated as the number of bases in a gain or loss state, divided by the total number of bases. Hereby excluding the difficult to assess centromeric (acen), variable-length (gvar) and tightly-constricted (stalk) regions. Likewise, the ploidy was derived from the weighted mean of the copy ratio of autosomal chromosome arms.

### Structural variants

Somatic structural variants (SVs) were detected with Manta (version 1.6) [46], DELLY(version 0.8.1) [47] and GRIDSS (version 2.7.2) [48]. First, we filtered SVs with a minimum of seven supporting reads and removed those with >90% reciprocal overlap with common (>1%) population variants retrieved from the NCBI repository (nstd166 [49], nstd186 [50]) and from DGV (version 2020-02-25) [51] accessed on 2021-03-11. Second, we merged SVs called by the three tools based on 50% reciprocal overlap and required detection by at least two tools. Third, we filtered on tumor allele fraction > 0.1 for all downstream analyses, except for dedicated analyses into the SV patterns of highly amplified regions (see below) since the allele fractions (AFs) of these SVs can be artificially low due to presence of many reference reads.

To identify whether SVs are likely repeat-mediated, we annotated them with repeats retrieved from UCSC table browser accessed on 2021-04-20 [29]. First, repeats were filtered by completeness (<50 base pairs (bp) of repeats left) and repeat class (LINE, SINE, LTR, Simple_repeat, Low_complexity, Retroposon) to prevent spurious annotations. To be considered repeat-mediated, we required both the start and end breakpoints within 100 bp of repeat elements of the same class.

### CN changes associated with interchromosomal breakpoints

To analyze whether interchromosomal breakpoints (CTX) are associated with chromosome level CN changes, we categorize CTXs as unbalanced, balanced or inconclusive. Hereto, we compared CN states before and after the breakpoint using windows that 1) extend across the full chromosome (arm) and 2) encompass the direct vicinity (5 Mbp flanking regions).

First, we defined windows relative to the chromosome and inferred their CN state and stability (see methods on CN alterations). Chromosome level windows extend from the 5’ telomere to the bp (“before”) and from the bp to the 3’ telomere (“after”). We required both before and after windows to be CN stable (>66%) and sufficiently large to assess (>10 Mbp), otherwise they were regarded as “inconclusive”. Second, we compared the CN of the before and after windows using a difference in CN call state or >0.2 cr l2fc as criterion for “unbalanced”. Third, we performed this analysis with the 5 Mbp before and after the CTX breakpoint to verify the chromosome level observations. For the final categorization of the CTX into unbalanced or balanced, we required the local CN changes to either match the chromosome level observation or be “inconclusive”, as such allowing for small CN changes around CTX breakpoints that are commonly observed. If the CTX could not be categorized on the chromosome level, chromosome arm-level windows were considered. These are defined similarly but using the nearest centromere instead of both telomeres, and >5 Mbp minimum size. Finally, CTXs within 5 Mbp distance of the telomeres were labeled as ’edge’ breakpoints.

In downstream analyses, we used the location of the breakpoint to the nearest telomere as the genomic interval of the CN change of unbalanced CTXs.

### Chromosome (arm) level CN alterations

To assess chromosome level CN alterations, we considered the most prevalent CN call state, the CN stability and the presence of unbalanced SVs. CN call states for chromosomes were identified based on the highest fraction of sequence. This allows for identification of numerical CN changes in the presence of focal CN changes, for example if a chromosome has 85% gain and 15% loss due to a large focal deletion, then the fraction in the “gain” call state will be selected as the chromosome-level CN state and further assessed for stability. As criteria for a chromosome level CN alteration, we required a >70% stable sequence of the gain, loss or LOH call state, and absence of an explanatory unbalanced translocation.

For chromosome-arm level alterations, the same approach was used but here we excluded the acen/gvar/stalk regions. In case CN alterations were identified on the full chromosome level, these take precedence over chromosome arm level alterations.

### Amplicons

Focal amplifications (amplicons) can arise through different mechanisms and exist in multiple forms, such as circular extrachromosomal DNA (ecDNA) constructs and linear homogeneous staining regions. ecDNAs tend to be smaller with an expected size between 10 kilobase pairs (kbp) and 10 megabase pairs (Mbp) and more highly amplified (> 8 copies) than local rearrangements [52]. To identify amplicons, we selected regions with high CN (>1.9 cr l2fc), corresponding to ∼7.5 copies to account for lower tumor cell fractions. We also applied the same threshold relative to the mean CN of the chromosome to account for the presence of chromosomal gains. For example, >2.4 cr l2fc would be the required CN in a 3n chromosome with mean 0.5 cr l2fc. Next, adjacent CN segments within 6 kbp were merged and we selected amplicons >50 kbp for downstream analysis.

Next, we analyzed whether separate amplicons are connected to each other by SVs (Table 2), which could indicate co-amplification on ecDNA. Hereto, amplicons were annotated by overlapping complex SVs (see next section) and we looked into whether low allele fraction SVs connect separate amplicons. For these SVs, the allele fractions are likely artificially low due to presence of many reference reads. Furthermore, we annotated amplicons with overlapping complex SVs and analyzed whether SVs have both breakpoints inside the amplicon (no loose ends).

### Identification of complex SVs

Complex SVs are characterized by clusters of breakpoints reflecting the repair of multiple simultaneous dsDNA breaks. To identify clusters of SVs, we used a graph-based approach considering SVs as vertices and drawing edges between SVs that have breakpoints within 5 Mbp. Connections between chromosomes can be formed by interchromosomal breakpoints which are represented as two vertices with an edge. From each tumor’s graph, we identified SV clusters by extracting the connected components, and subsequently categorized them into different types of (complex) events based on a combination of SV and CN features.

After identifying clusters of SVs, they were categorized into different complex and non-complex classes: ecDNA, pair of SVs, reciprocal translocation, chromoplexy, chromothripsis and ’complex other’. See below the criteria used for each class, and also the order of categorization is of importance:

1) ecDNA/amplicon : SV breakpoint within 1 kbp of an amplicon.

We first categorized this class to distinguish potential ecDNA from chromoanagenesis events. Although we could not definitively establish with short-read WGS whether the complex SV clusters represent circular ecDNA constructs, we selected the amplicon criteria to optimize for this [15] and detected closed chains in graphs for many clusters as expected [52], but we did not require this as criterium given that SV calling is challenging in highly amplified regions. Instead, we analyzed whether SVs had both breakpoints in an amplicon and observed that there were little to no “loose ends” for most ecDNA/amplicon clusters.

2) (non-complex) SV pair: two nearby SVs or one CTX consisting of two breakpoints and a nearby SV.

Although we do not regard two nearby SVs as “complex events”, also small clusters of two breakpoints were extracted to allow for detection of chained interchromosomal events having just two breakpoints on a chromosome.

3) (non-complex) Reciprocal translocation: four CTXs connecting two chromosomes, with <2 Mbp distance between the CTX breakpoints indicating a simple rearrangement. Although the presence of small intrachromosomal SVs is allowed, the SVs should form a closed chain and footprints remain <20 Mbp.

4) Chromoplexy: a closed chain of SVs connecting two or more chromosomes. No unbalanced CTXs are allowed and all chromosomes have at least one balanced or edge CTX.

5) Chromothripsis: high density of SVs, namely at least 10 overlapping SVs or 10 CTXs between a chromosome pair. As a proxy for CN oscillations, we merged all adjacent CN segments with the same call state, and required at least 10 CN state switches within a footprint. Finally, the distribution of SV types should not be significantly different from an equal distribution (chi-squared test p-value > 0.05) indicating randomness of fragment joining.

6) Complex other: SV clusters not satisfying the criteria outlined above.

### Genomic regions recurrently affected by SVs

Recurrently altered regions were identified using a peak-calling approach based on SV breakpoints. For a conservative measure of recurrence, we counted the number of distinct tumors with a breakpoint in the region, such as to not bias for certain complex SV classes that inherently have many bps. First, we overlapped all SV breakpoints using 1 Mbp flanking windows and partitioned the genome into regions based on the number of distinct tumors. Next, we mapped each breakpoint to the highest hotspot it overlaps with, having the most distinct tumors. In case of a tie, we selected the region with the highest fraction of overlap with the breakpoint. After this mapping, we re-assessed the number of distinct tumors contributing to each region. Finally, we annotated the genomic regions with cancer genes and assessed the fraction of SVs from each cancer type and complex SV class.

Complex SV hotspots were defined as recurrently altered regions with complex SV breakpoints from three or more tumors.

### CN alterations due to complex SVs

To associate chromosomal alterations with complex SVs, we combined overlap with CN segments and unbalanced translocations. First, we inferred focal CNs relative to the chromosome average and merge CN segments with the same state, allowing for up to 5 Mbp gaps to ignore interruption by small fluctuations. Then, these merged CNs were matched to complex SVs if they overlapped at least 5 Mbp, or at least half their size for smaller segments. Second, we added the CN changes due to the unbalanced translocations part of the SV cluster, as defined previously. This allowed us to conservatively infer a set of CN changes likely due to a complex SV event, and ignore potential co-occurring chromosomal changes such as aneuploidies.

### Alterations affecting cancer driver genes and chromosomes

To assess whether a gene is altered, we combined SNV/indels, CNs and SVs and filtered alterations that could affect gene function. The alteration types reported in Table 4 and Figure 3 are based on the following:

- SNV: SNV or indel predicted by VEP to have moderate or high impact
- SV: breakpoint inside the gene body, or breakpoint within 1 Mbp of the gene body together with a CN change
- Complex: the SV breakpoint is part of a complex SV
- CNA: high-impact CN change by itself namely homozygous loss (-1 cr l2fc) or amplification (>1.9 cr l2fc)

Chromosomal alteration types reported in Figure 3:

- Complex: CN alterations due to complex SVs
- SV: CN changes associated with interchromosomal breakpoints
- CNA: Chromosome (arm) level CN alterations

Only genomic regions >5 Mbp are included in the figure and downstream analyses.

To identify alterations relevant for tumor biology, we focused on pediatric cancer genes (Table 7) and more specifically on previously established cancer-type specific associations with genes and chromosomal alterations. The following definitions were used: Cancer driver genes: genes in which alterations confer a selective advantage to the tumor (Table 8). Unfavorable chromosomal alterations: chromosomal alterations that have been associated with poor outcome or high risk (Table 9).

Complex SVs were annotated with their ‘effect’ based on whether they affect cancer driver genes (*driver)* or result in unfavorable chromosomal alterations (*chrom_alt)*.

### Statistical tests

Fisher’s exact test was used to compare two groups and assess enrichments.

### Fluorescence in situ hybridization

Fluorescence in situ hybridization analysis was performed using Vysis LSI FOXO1 (13q14) Dual Color, Break Apart Rearrangement Probe 30-231023.

## Tables

**Table 1: Complex SV clusters**

Complex SV clusters annotated with characteristics supporting their categorization as a certain complex class and to assess their pathogenicity, e.g. affected cancer driver genes and summary of copy number changes.

**Table 2: Amplicons**

Amplicons annotated with genes and (complex) SVs for assessment of physical linkage.

**Table 3: Genome-wide SV hotspots**

Genomic regions recurrently altered by SVs in at least three tumors, annotated with cancer genes. Complex SV hotspots have complex SV breakpoints from at least three tumors.

**Table 4: Alterations in cancer driver genes**

Overview of SNVs, indels, CNs and SVs identified in cancer driver genes (see Table 8).

**Table 5: Copy number changes due to complex SVs**

Copy number changes associated with complex SVs, both overlapping local changes and unbalanced translocations.

**Table 6: Complete cohort overview**

Patient information related to diagnosis, and if applicable treatment and clinical events, as well as identifiers of the WGS data and quality assessment.

**Table 7: Pediatric cancer gene panel**

**Table 8: Cancer driver genes per cancer type**

**Table 9: Unfavorable chromosomal alterations per cancer type**

## Declarations

### Availability of data and materials

All underlying data is included in the supplementary materials. The WGS datasets supporting the conclusions of this article are available in the EGA repository under study EGAS00001007565. Data from organoid sample PMOBM000ABW is available in EGAD00001008466. Code is available through https://github.com/princessmaximacenter/structuralvariation/ (tag v2.0), released on 2023-09-18.

### Ethics approval and consent to participate

Informed consent has been obtained for all subjects involved in this study through the Máxima biobank informed consent procedure and corresponding protocol. The Máxima biobank protocol has been approved by the Medical Ethics Committee of the Erasmus Medical Center in Rotterdam, The Netherlands, under reference number MEC-2016-739. Approval for use of the subject’s data within the context of this study has been granted by the Máxima biobank and data access committee, biobank request nr. PMCLAB2018.017.

### Competing interests

The authors declare no competing interests.

### Funding

We gratefully acknowledge financial support provided by the Foundation Children Cancer Free (KiKa core funding) and Adessium Foundation. The funders had no role in the design of the study, as well as the collection, analysis and interpretation of the data or writing.

### Author contributions

Conceptualisation: I.A.E.M.vB., J.Y.H.K., P.K., F.C.P.H. and B.B.J.T.; supervision: P.K., J.Y.H.K, F.C.P.H. and B.B.J.T.; funding acquisition: P.K.; methodology: I.A.E.M.vB., J.Y.H.K. and P.K.; investigation and visualization I.A.E.M.vB.; software: I.A.E.M.vB., S.B., A.J., E.T.P.V., M.S., S.dV., J.B.H. and H.H.D.K.; resources: M.vT, N.S.W.; data curation: I.A.E.M.vB., S.B., A.J., E.T.P.V., M.S., S.dV., J.B.H., H.H.D.K; writing - original draft: I.A.E.M.vB., J.Y.H.K. and P.K.; writing - review & editing: I.A.E.M.vB., J.Y.H.K., P.K., M.T.M., N.S.W., J.D., M.M.vdH.E., J.H.M.M, J.J.M., W.C.P., and B.B.J.T.

## Acknowledgments

We acknowledge the willingness of all children and their families to donate their material and data for our research. We also want to thank the contributions from all pediatric oncologists, research nurses, lab technicians and biobank committee members for obtaining patient consent, sample collection and processing as well as general logistics. We acknowledge the fruitful discussions and feedback with fellow lab members as well as Ivo Griffioen.

## Notes

### Competing Interest Statement

The authors have declared no competing interest.

