## Supplementary Figures for "Complex structural variation is prevalent and highly pathogenic in pediatric solid tumors"

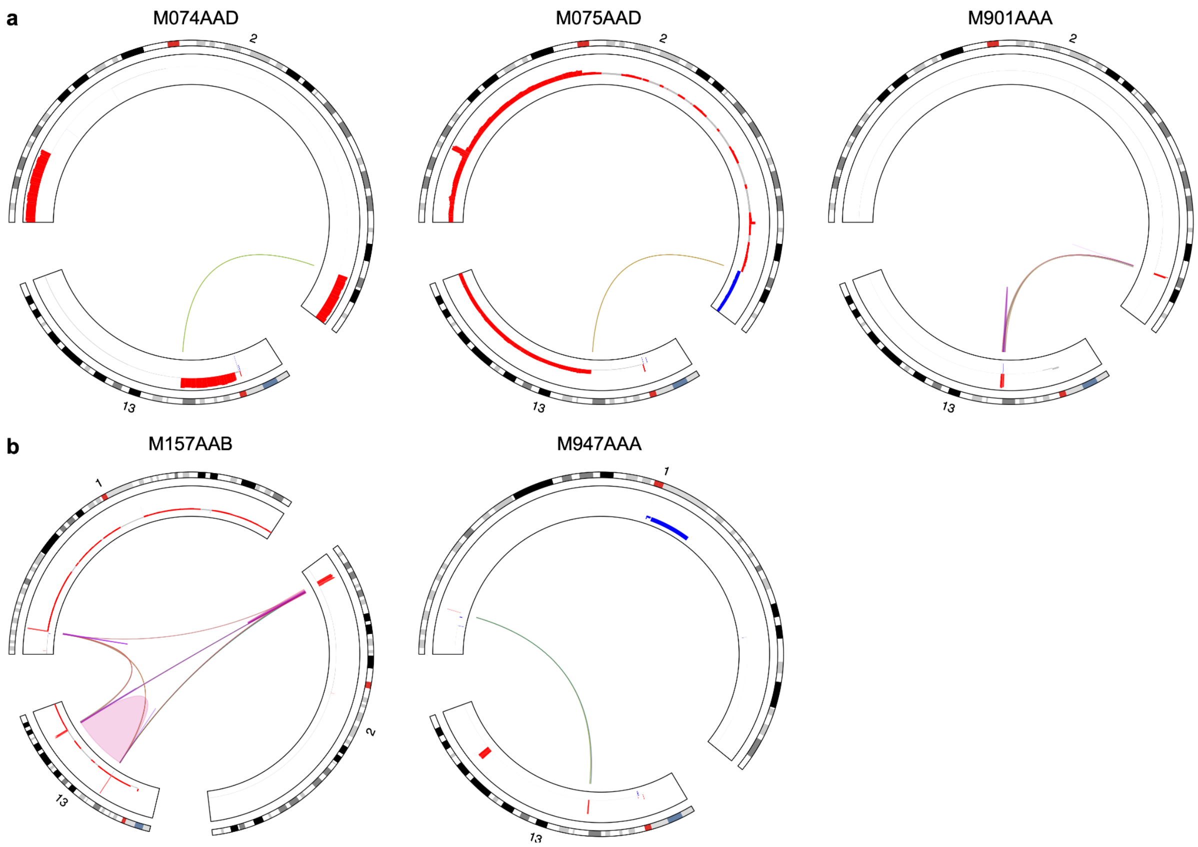


**Figure S1: Distinct underlying rearrangements of *PAX3/7::FOXO1* fusion genes**

Circos plots of the complex SVs underlying driver gene fusions. In all tumors we identified interchromosomal breakpoints (CTX) connecting *PAX3* or *PAX7* to *FOXO1* but they are part of quite different genomic rearrangements.

**a** *PAX3::FOXO1* fusions: M074AAD has a reciprocal translocation with terminal chr2q gain and interstitial chr13q gain. M075AAD has a reciprocal translocation with terminal chr2q loss and chr13q terminal gain and copy number data indicated the tumor is likely tetraploid. For M901AAA the fusion arises from chromoplexy: there are eight CTX between chr2-chr13 and it is balanced at the chromosomal level with focal gains adjacent to the breakpoint (1.1 copy ratio log2 fold change (cr l2fc)).

**b** *PAX7::FOXO1* fusions: In both M157AAB and M947AAA we identified ecDNA-type complex SVs with CTX connecting focal amplifications overlapping *PAX7* and *FOXO1*. Only the parts of the genes that are expected to be included in the fusion product are amplified. Namely the 5' end of PAX7: for M947AAA 2.0 cr l2fc 67% of the gene, and for M157AAB 3.7 cr l2fc (67%). And the 3' end of FOXO1: for M947AAA 2.0 cr l2fc (72%) and for M157AAB 3.8 cr l2fc (47%). In the case of M157AAB, we identified breakpoints connecting the ecDNA with the fusion gene to a *MYCN* amplification (3.0 cr l2fc).

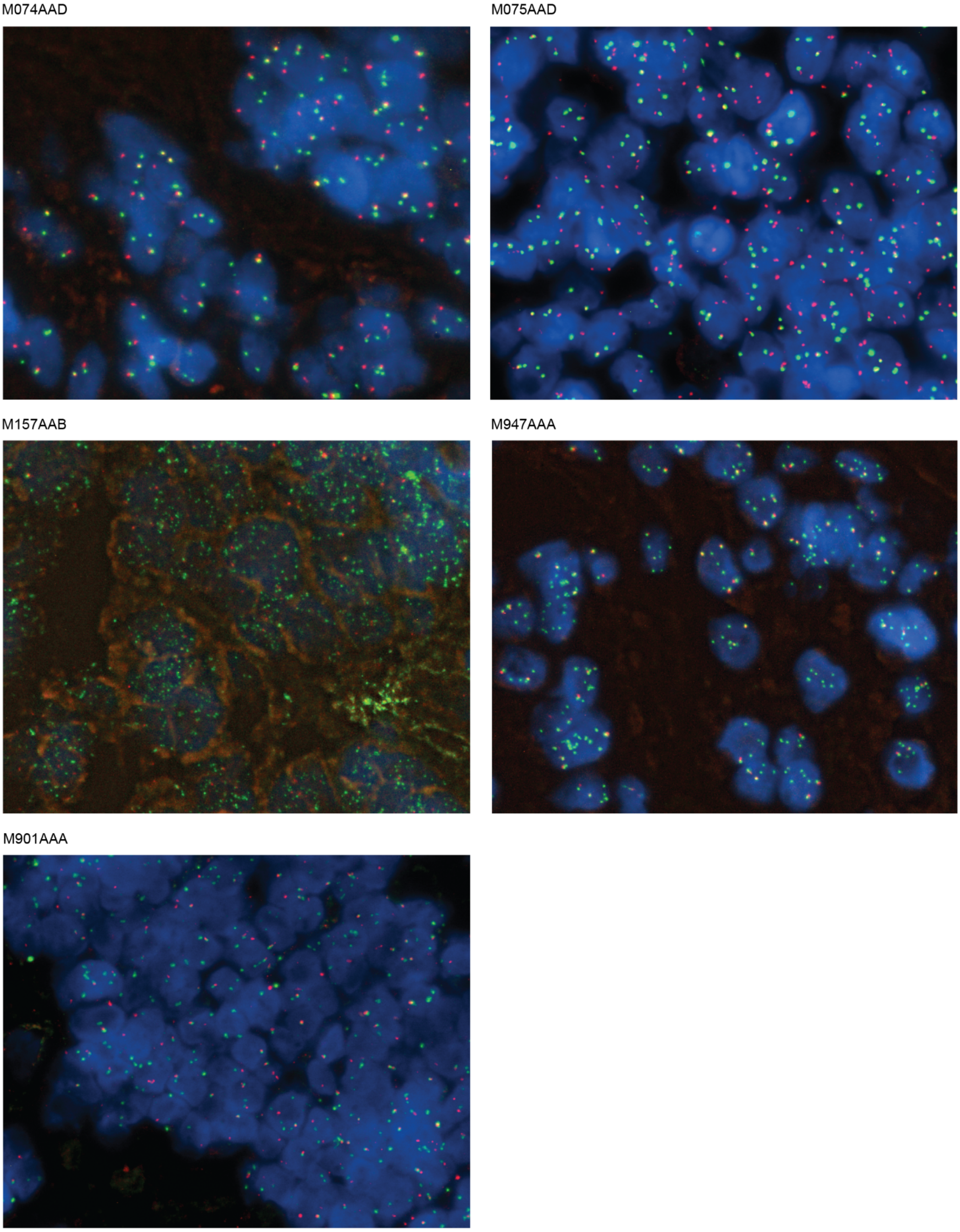


**Figure S2: *FOXO1* break-apart fluorescence *in situ* hybridization**
Fluorescence *in situ* hybridization using *FOXO1* break-apart probes supports the observed differences in WGS data**.** For M074AAD and M075AAD with reciprocal translocations, a clear break-apart signal could be observed, as well as break-apart and gain for M901AAA with chromoplexy. In contrast, for M157AAB and M947AAA, we instead observed many separate green signals, as expected for fusion genes residing on ecDNA constructs (Figure S1) [1].

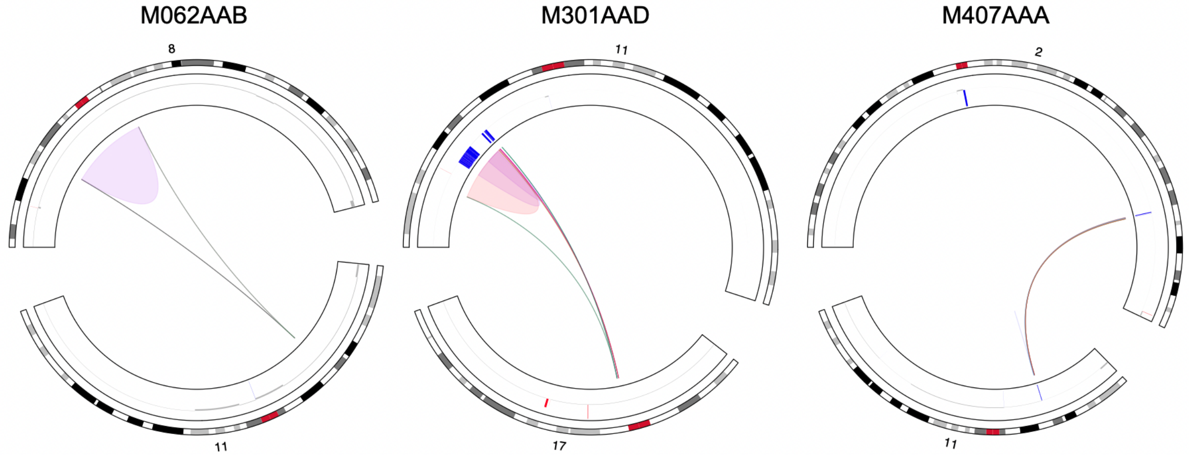


**Figure S3: Chromoplexy with breakpoints in WT1 in three Wilms tumors**

Circos plots of the chromoplexy events identified in tumors from patients M062AAB, M301AAD and M407AAA (left to right). These complex SVs have breakpoints in *WT1* and result in rearranged chromosomes, leading to substantial disruption of the gene.
For tumor M062AAB, the complex SV is fully copy number balanced whilst there are focal deletions at the breakpoints for M301AAD and M407AAA. Not visible in circos plot: M407AAA has eight interchomosomal breakpoints (Table 1).

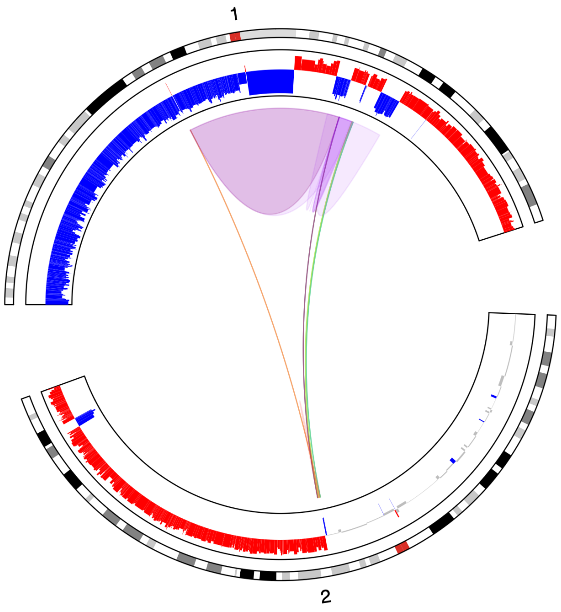


**Figure S4: Complex SV identified in hepatoblastoma tumor from patient M147AAD**

Associated with unfavorable chromosomal alterations 1p loss, 1q gain and 2q gain [11,12].

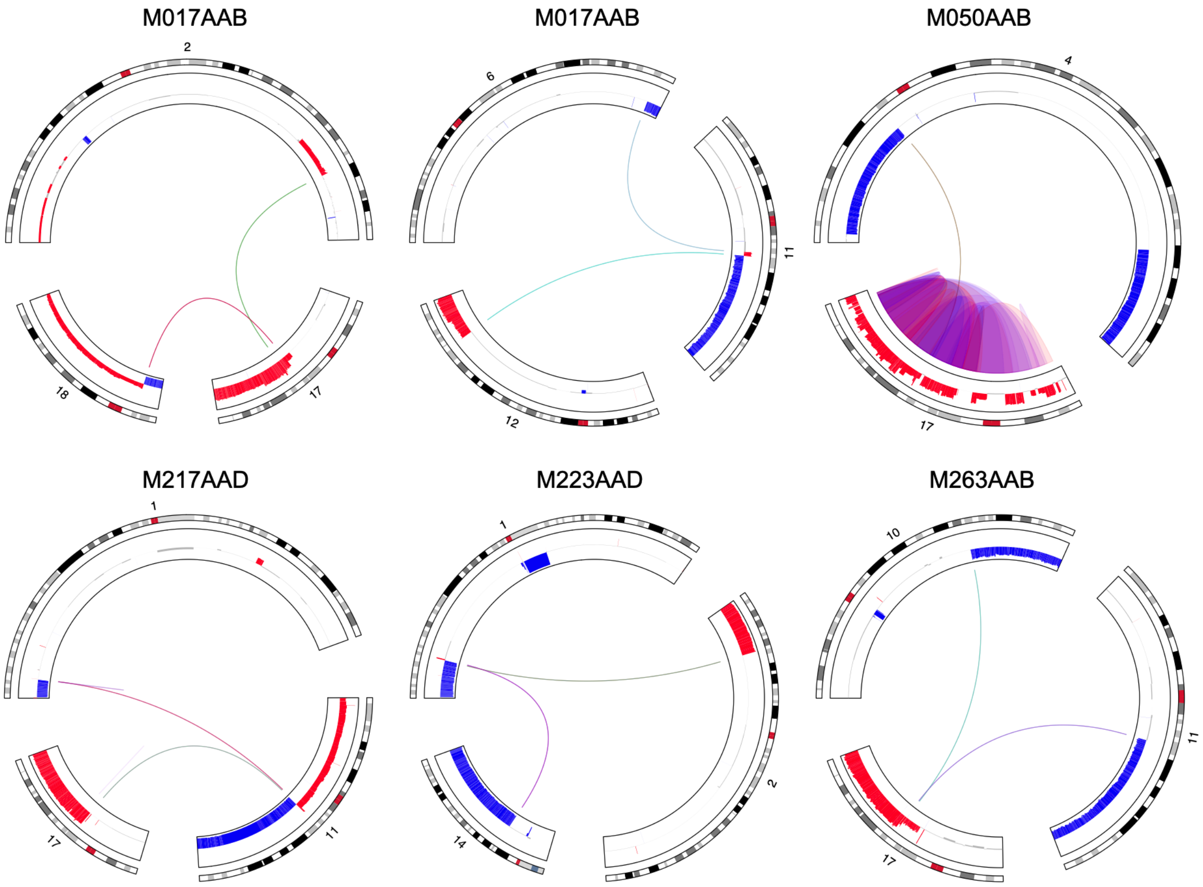


**Figure S5: Complex SVs in neuroblastoma that result in unfavorable chromosomal alterations**

Circos plots displaying the complex SVs and the resulting unfavorable chromosomal alterations for patients (top left to bottom right): M017AAB chr17q gain and chr11q loss; M050AAB additional chromothriptic copy of chr17 (note that complex sv cluster contains unrelated interchromosomal breakpoint to chr4); M217AAD chr1p loss, chr11q loss and chr17q gain; M223AAD chr1p loss; M263AAB chr10q loss, chr11q loss, chr17q gain [2–4].

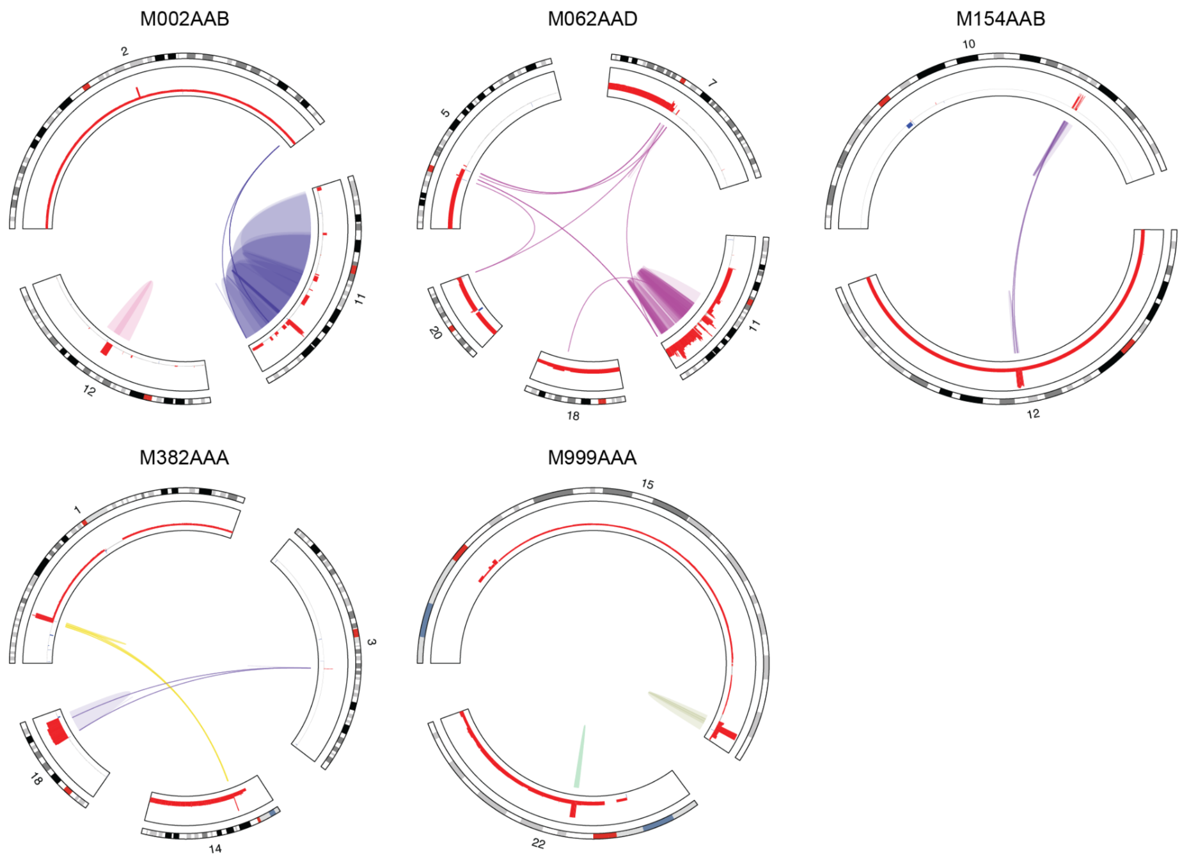


**Figure S6: ecDNA/amplicon-type complex SVs with cancer driver genes in RMS**

Circos plots displaying ecDNA/amplicon-type complex SVs identified in fusion-negative RMS that result in oncogene amplification, coloured by event. From top left to bottom right:

M002AAB has two ecDNA/amplicons, one resulting in amplification of *MDM2* on chr12 (4.2 copy ratio log2 fold change (cr l2fc)) and another of *YAP1* (7.9 cr l2fc) on chr11, but we did not find evidence that they are connected (Table 2). The chr11 complex SV spans a large region, nevertheless it involves 2.5 megabase pairs (Mbp) amplified bases so could represent an ecDNA arising from chromothripsis. The role of the interchromosomal breakpoints (CTX) is unclear and could be part of a different alteration.

M062AAD has an ecDNA/amplicon containing *YAP1* (2.0 cr l2fc). This complex SV could represent an ecDNA resulting from chromothripsis, but could also reflect an homogeneous staining region given the lower amplification amplitude and large footprint spanning over 30 Mbp. The CTXes likely reflect a different event with unbalanced translocations, possibly involving another homolog.

M154AAB has an ecDNA/amplicon with *MDM2* (3.4 cr l2fc) together with a locus on chr10. The complex SV includes only breakpoints connecting the amplicons and also the high copy ratio is consistent with ecDNA.

M382AAA has an ecDNA/amplicon with *MYCL* (5.3 cr l2fc) on chr1 connected to chr14, and also carries a complex SV connecting amplicons on chr3 and chr18. The number of amplified bases of the chr18 locus is too large to likely be ecDNA (20.9 Mbp).

M999AAA has two ecDNA/amplicons with *IGF1R* (5.3 cr l2fc) on chr15, and on chr22 containing the pediatric cancer genes *CRKL* and *LZTR1*. We did not find evidence that they are connected.

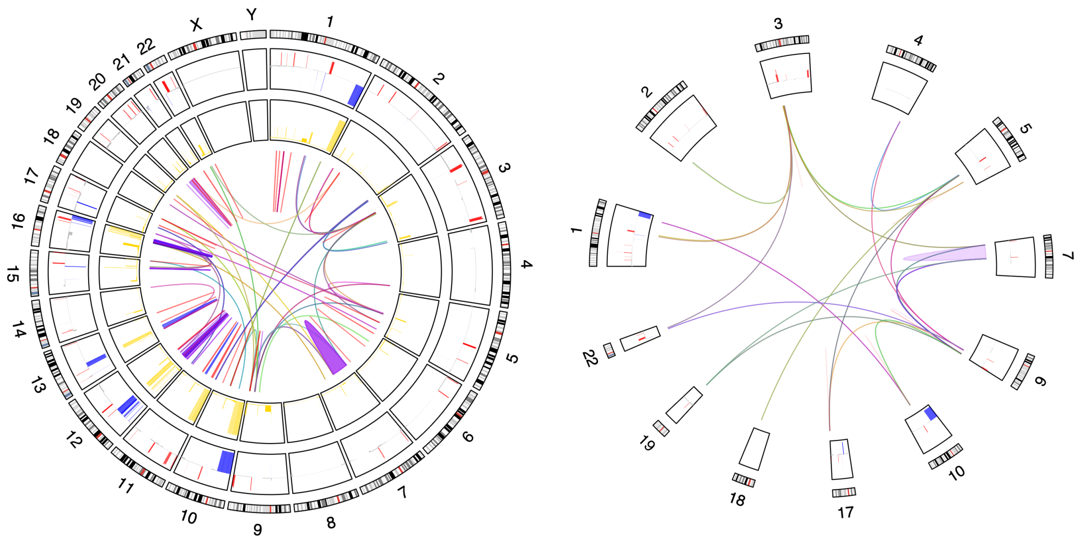


**Figure S7: Complex SVs identified in tumor from M365AAD**

The tumor from patient M365AAD shows a remarkable genome-wide alteration pattern of interconnected focal gains. The gains are of similar amplitude (~0.5 cr l2fc) and often connected by balanced interchromosomal breakpoints (~0.25 tumor allele fraction). Varying in size from 15 kilobase pairs (kbp) to 7.3 megabase pairs (Mbp), the focal gains are too large to be templated insertions (<1 kbp). Although the focal gains do not meet the thresholds for amplification, this pattern is reminiscent of ecDNA so we hypothesized that it could be explained by subclonal ecDNAs. This could indicate high intratumor heterogeneity and/or ongoing genomic instability.

The largest complex SV is comprised of focal gains from 12 chromosomes with cr l2fc 0.13-0.37 (median 0.24 cr l2fc). A construct like this could arise from ecDNAs that cluster together and then get rearranged [25].

Furthermore, M365AAD is one of two tumors with a *MYOD1* L122R mutation (0.95 tumor AF), which is associated with dismal outcome. But it is the only tumor displaying this pattern of connected focal gains. It appears to be diploid with a low fraction of genome altered by copy numbers (7.5%) of which half due to complex SVs, whilst other fusion-negative RMS tend to be hyperdiploid.

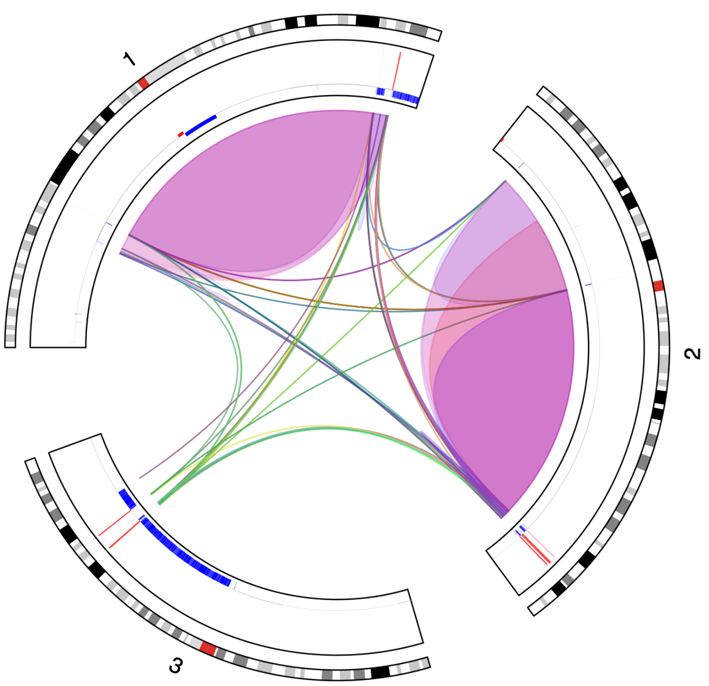


**Figure S8: ecDNA resulting in *PAX3::WWRT1* fusion gene in the tumor of M911AAA**

Circos plot of the complex event identified for patient M911AAA that connects amplified loci on chromosomes 1, 2 and 3 (~4.5 cr l2fc) and gives rise to the rare fusion gene *PAX3::WWRT1* with very high breakpoint allele fractions (~0.9)

The observed pattern is consistent with the hypothesized translocation-excision-deletion-amplification mechanism underlying *PAX7::FOXO1* fusion genes [5] and similar to patterns we observed in *PAX7::FOXO1* fusion-positive tumors in our cohort (Figures S1 and S2).

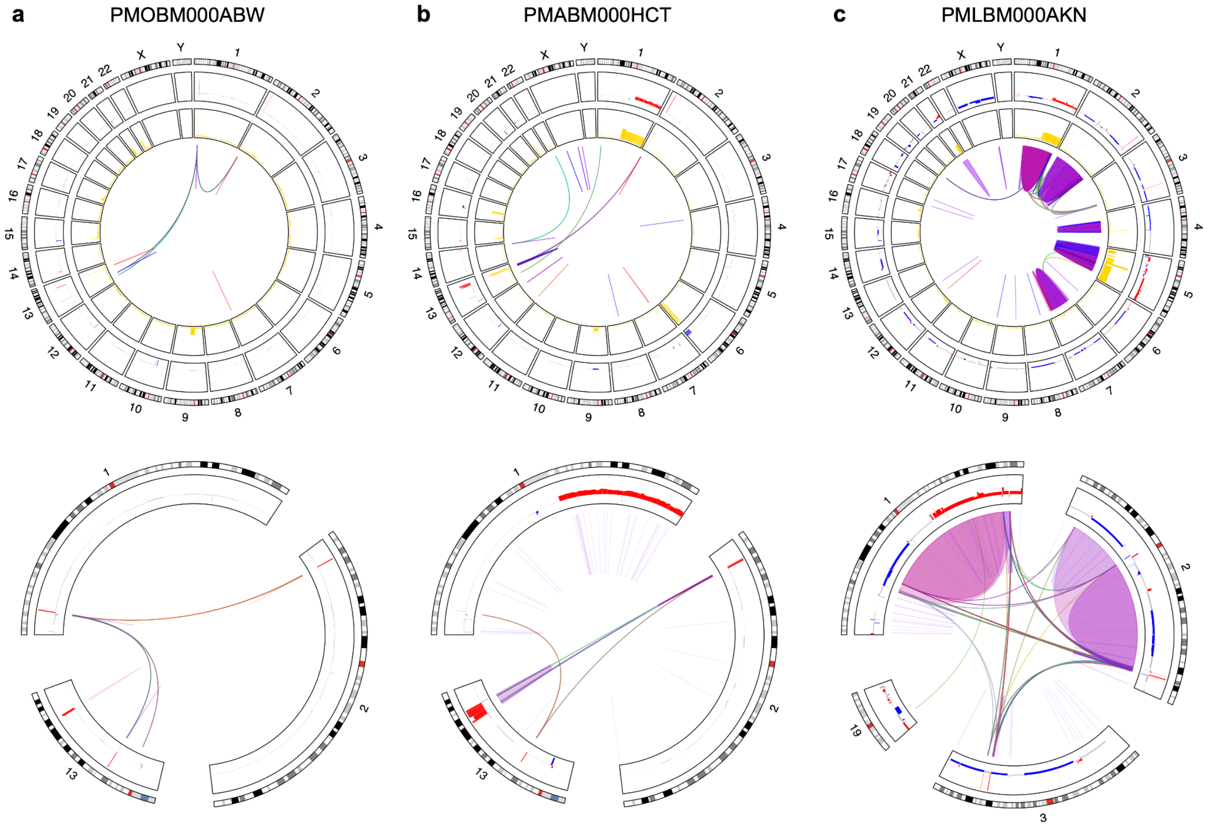


**Figure S9: *PAX3/7::FOXO1* fusions identified in relapse and organoid samples**

Circos plots of additional samples from three patients with RMS (top row) and the complex events we identified that give rise to gene fusions (bottom row).

**a** Organoid sample (PMOBM000ABW [6]) from M157AAB contains the same breakpoints as the primary tumor underlying the fusion gene *PAX7::FOXO1* connected to *MYCN* amplification.

**b** Relapse sample of M947AAA (232 days PMABM000HCT) contains the same breakpoints as primary tumor underlying the fusion gene *PAX7::FOXO1*, and additionally acquired *MYCN* amplification in the same ecDNA.

**c** Relapse sample of M911AAA (430 days PMLBM000AKN) contains the same breakpoints as the primary tumor underlying the fusion gene *PAX3::WWTR1*.
